# Bat power-metabolic profiling of the Egyptian fruit bat *Rousettus aegyptiacus* reveals distinctive cardiac adaptations

**DOI:** 10.1101/2025.04.16.649087

**Authors:** Anja Karlstaedt, Fenn Cullen, Rosie Drinkwater, Kyoungmin Kim, Megan Young, Stephen J. Rossiter, Dunja Aksentijevic

## Abstract

**Aim:** The present study aimed to elucidate which pathways contribute to cardiometabolic adaptation in Egyptian fruit bats.

**Methods:** Utilising cardiac tissues from Egyptian fruit bats (*Rousettus aegyptiacus*) and C57BL/6J mice, we combined liquid chromatography-mass spectrometry metabolic profiling, non-targeted ^1^H NMR spectroscopy, and *in silico*computational modelling using the genome-scale mammalian network CardioNet.

**Main Findings:** Our analyses revealed that bat hearts exhibit a distinct metabolic profile characterised by depleted glycogen reserves and increased reliance on lipid oxidation to meet energy demands. Notably, bat hearts displayed elevated fluxes in oxidative phosphorylation, β-oxidation of long-chain fatty acids, and the Krebs cycle, alongside reduced amino acid catabolism. These findings suggest that bats have evolved unique metabolic strategies to support the high-energy demands of flight, maintaining cardiac function without succumbing to pathological remodelling.

**Conclusions:** This study provides the first comprehensive insight into the metabolic adaptations in the cardiac tissue of a bat species, contributing to our understanding of how these mammals endure extreme physiological stresses. Integrating metabolomics with computational modelling offers a powerful approach to studying metabolic adaptations in non-model species, potentially informing therapeutic strategies for human cardiac conditions.

## 1. INTRODUCTION

Bats are exceptional among mammals for sustaining the extreme metabolic demands of powered flight while achieving lifespans far exceeding those of similar-sized non-flying species^1^. Bats also show adaptations in mechanisms underlying DNA repair and inflammatory responses^2-5^, and studies conducted on multiple species across several decades have highlighted their capacity for extreme cardiovascular performance under fluctuating energetic demands and resistance to oxidative stress^2,3,6-14^.

Flight drives rapid and reversible increases in heart rate and cardiac workload; for example, free-flying bats can exceed 800 beats per minute yet quickly return to low resting rates or even enter torpor under energy-saving conditions^3,12,15^. When resting, many bat species roost in caves or enclosed environments in which microclimatic conditions can be characterized by elevated CO□ levels and reduced air exchange^16^, thus imposing additional respiratory and metabolic challenges. This combination of energetic extremes alongside remarkable longevity makes the bat heart a fascinating model for uncovering metabolic strategies beyond those of canonical mammalian physiology.

Several earlier studies have suggested that bat hearts possess unique molecular and physiological traits. For example, comparative genomics reveals alternative splicing of *TNNI3*(cardiac troponin I) isoforms that may facilitate rapid diastolic relaxation at high heart rates without excessive adrenergic stimulation^17^. Mitochondrial studies indicate lower reactive oxygen species production despite high lifetime oxygen fluxes, potentially explaining bats’ resilience to age-related cardiovascular decline^1,6^. Furthermore, frugivorous and nectarivorous bats exhibit dramatic postprandial hyperglycaemia, and stable isotope studies in nectar bats show systemic metabolic switching from lipid oxidation when fasted to carbohydrate oxidation after feeding^18-22^. Yet how these features integrate within the bat myocardium to support high energetic demand without pathological remodelling remains unresolved.

Here, we combine high-resolution metabolomics with *in silico*flux balance analysis to compare the cardiac metabolic networks of the Egyptian fruit bat (*Rousettus aegyptiacus*) and the laboratory mouse. The rationale for our metabolomic and computational approach is to identify systems-level metabolic features that may underpin the extraordinary cardiac performance of the Egyptian fruit bat heart. We hypothesise that the bat heart is metabolically reprogrammed towards enhanced mitochondrial oxidative pathways, particularly long-chain fatty acid β-oxidation and ketone metabolism while maintaining minimal cytosolic energy reserves, enabling sustained ATP production and redox balance under extreme energetic stress. We predict that myocardial metabolomics and flux balance analysis will reveal elevated oxidative phosphorylation metabolic fluxes in bats relative to mice, accompanied by lower cytosolic energy reserve metabolites.

## 2. MATERIAL AND METHODS

### 2.1. Animals

Cardiac tissue from Egyptian fruit bat (*R. aegyptiacus*, n=4) was obtained opportunistically during Copenhagen Zoo (Denmark) population management procedures (study methodological summary shown in Figure 1A). Animals were euthanised by trained veterinarians in line with management decisions in the zoo to ensure the best welfare of the captive colony. The feeding status of individual bats at the time of sampling could not be controlled or directly assessed. 20-week-old C57/Bl6J mice (n=4, male) were purchased from Charles River, UK (Figure 1A). Mice were housed in individually ventilated cages and maintained under controlled temperature (22 ± 2 °C) on a 12:12 h light-dark cycle with access to a standard chow diet (irradiated PicoLab® Mouse Diet 20 EXT, 5R58) and water *ad libitum*. Mice were sacrificed by phenobarbital I.P. injection, and hearts were rapidly excised. Mouse experiments were conducted by the UK Home Office Animals (Scientific Procedures) Act, 1986.

**Figure 1.**
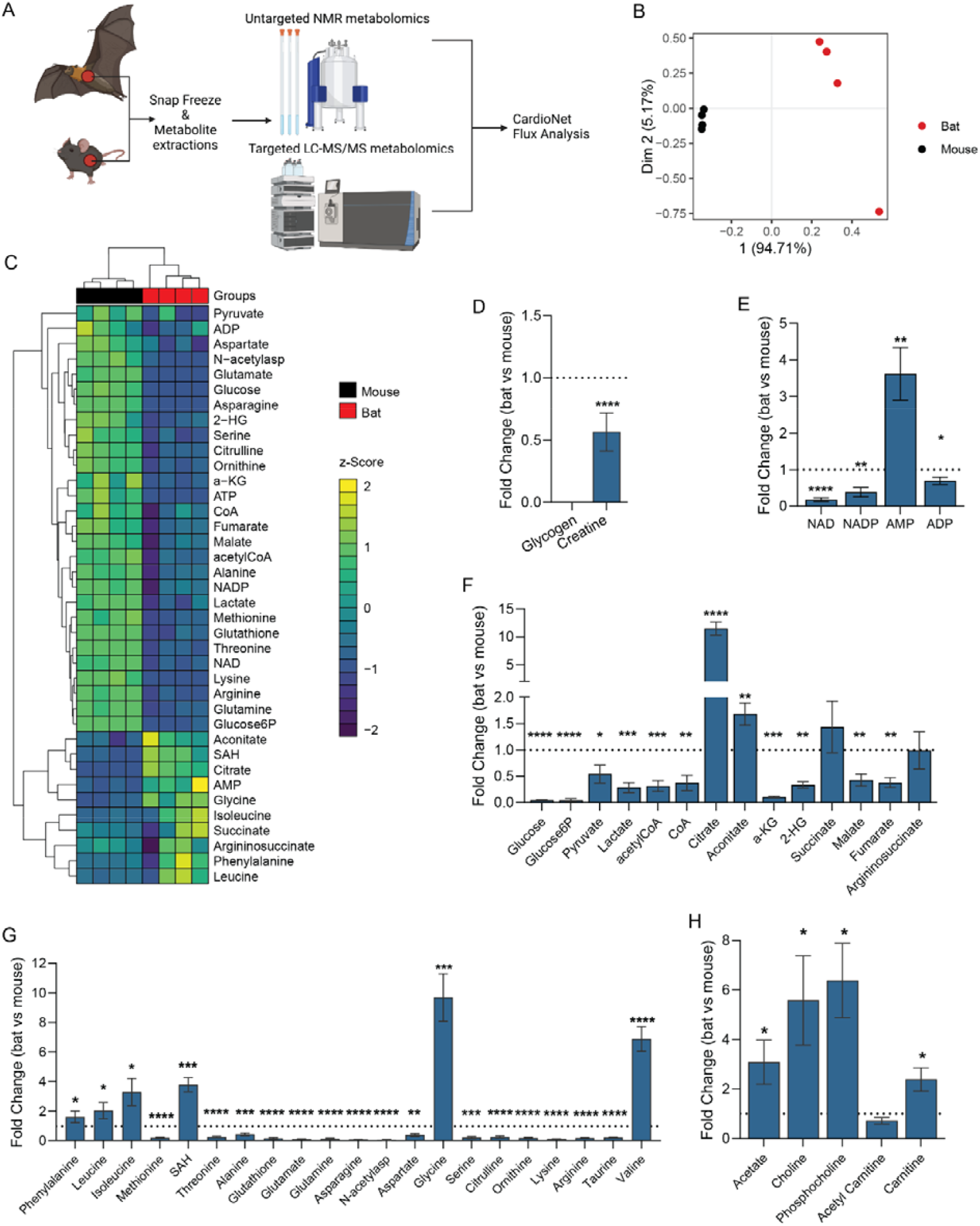
Cardiac metabolism varies between Egyptian fruit bats and C57BL/6J mice. **(A**) Schematic summary of the study protocol. n=4 bats (n=2 males, forearm length=91.60±2.50 mm, n=2 females, forearm length=87.90±1.85 mm) and n=4 C57/BL6J mice (Charles River, UK, male, 6 weeks). (**B-C**) Partial Least Squares Discriminant Analysis (PLS-DA) (**B**) and heatmap of unsupervised hierarchical cluster analysis (**C**) of the cardiac metabolite concentrations determined by LC-MS and high-resolution ^1^H nuclear magnetic resonance spectroscopy showing distinct profile of bat and C57/BL6J mouse. (**D-H**) Fold change differences of myocardial intermediates within major metabolic pathways (glucose metabolism, lipid metabolism, Krebs cycle), redox equivalents and energy-providing substrates. Fold changes are expressed as comparisons between bats and C57/BL6J mouse. The normality of data distribution was examined using Shapiro–Wilk’s normality test. Comparison between two groups was performed by Student’s t-test (Gaussian data distribution) or Mann-Whitney U test when data was not normally distributed, *p<0.05, **p<0.01, ***p<0.001, ****p<0.0001.

### 2.2. Heart tissue collection

Dissected hearts were stabilised in RNA later and cryogens (liquid nitrogen, dry ice). Tissue was stored at - 80°C until analysis. Metabolites were extracted from frozen tissue as previously described^3,4^.

### 2.3. Metabolomics

#### 2.3.1. ^1^H Untargeted NMR spectroscopy

Frozen, pulverised cardiac tissue samples (∼45 mg) were subject to methanol/water/chloroform phase extraction and analysed using ^1^H NMR high-resolution spectroscopy as previously described^23,24. 1^H NMR spectra were acquired using a vertical-bore, ultra-shielded Bruker 14.1. Tesla (600 MHz) spectrometer with a BBO probe at 298K using the Bruker noesygppr1d pulse sequence. Acquisition parameters were 128 scans, 4 dummy scans, and 20.8 ppm sweep width, acquisition time of 2.6s, pre-scan delay of 4s, 90° flip angle, and experiment duration of 14.4 minutes per sample. TopSpin (version 4.0.5) software was used for data acquisition and metabolite quantification. FIDs were multiplied by a line broadening factor of 0.3 Hz and Fourier-transformed, phase and automatic baseline-correction were applied. Chemical shifts were referenced to the TSP signal. Metabolite peaks of interest were initially integrated automatically using a pre-written integration region text file and then manually adjusted where required. Assignment of metabolites to their respective peaks based on previously obtained in-house data, confirmed by chemical shift and using Chenomx NMR Profiler Version 8.1 (Chenomx, Canada). Peak areas were normalised to the total metabolite peak area. Glycogen was quantified using ^1^H magnetic resonance, measuring the concentration of glucose monomers from normalised peaks. Being a large macromolecule, with possible differences in the mobility of glycosyl units, glycogen has been reported to be fully visible by MRS. The modified dual-phase Folch extraction method, used for separating aqueous and lipid metabolites, was not optimised for extracting glycogen. However, all samples underwent the same extraction procedure, allowing for comparison between groups ^23^.

#### 2.3.2. Targeted LC-MS/MS

Frozen pulverised cardiac tissue samples (∼15 mg) were extracted using a buffer (50% Methanol, 30% Acetonitrile, 20% ultrapure water, and 50 ng/ml 1 mM HEPES solution) for LC-MS analysis. LC-MS analysis was performed by the metabolic flux analysis facility of the Barts Faculty of Medicine and Dentistry using a Q Exactive Quadrupole-Orbitrap mass spectrometer coupled to a Vanquish UHPLC system (Thermo Fisher Scientific). The liquid chromatography system was fitted with a Sequant ZIC-pHILIC column (150 mm × 2.1 mm) and guard column (20 mm × 2.1 mm) from Merck Millipore (Germany), and the temperature was maintained at 35°C.□The sample (3μL) was separated at a flow rate of 0.1 mL/min. The mobile phase comprised 10 mM ammonium bicarbonate and 0.15% ammonium hydroxide in water (solvent A), and acetonitrile (solvent B). A linear gradient was applied by increasing the concentration of A from 20 to 80% within 22 min and then maintained for 7 minutes.□The mass spectrometer was operated in full MS and polarity switching mode, in the range of 70-1000m/z and resolution 70000. Major ESI source settings: spray voltage 3.5kv, capillary temperature 275°C, sheath gas 35, auxiliary gas 5, AGC target 3e6, and maximum injection time 200ms.□□The acquired spectra were analysed using XCalibur Qual Browser and XCalibur Quan Browser software (Thermo Scientific) for the targeted MS analysis.

### 2.4. CardioNet-based metabolic flux analysis

*In silico*simulations were conducted using the publicly available metabolic network of the mammalian heart metabolism, CardioNet^25-27^. Mathematical modelling has previously been used to study the dynamics of cardiac metabolism in response to stress. Metabolite abundances from NMR and LC-MS/MS analysis were integrated as constraints for metabolite pools. The objective was to maximize ATP hydrolysis as a reflection of cardiac contraction (vATPase) that satisfies the metabolic constraints. We determined flux distributions (*v*_*m*_) and estimated flux rate changes (v_FC_) as described previously^24,25,28,29^. We simulated rapid flight, night, and roost activity states using published telemetry heart rate data for the neotropical frugivorous bat *Uroderma bilobatum*^15^. To our knowledge, detailed continuous telemetry datasets are not currently available for *R. aegyptiacus*. We therefore used *U. bilobatum*as an ecologically comparable fruit bat to estimate cardiac ATP demands under different behavioral states. Importantly, the underlying cardiac metabolic data (NMR, LC-MS/MS) used to constrain the model fluxes came directly from *R. aegyptiacus*heart tissue.

Side constraints and parameters for FBA are provided in the Supplementary Material to enable reproducibility and adaptation with new telemetry data as it becomes available. The demand for ATP per heartbeat was based on the transient ATP increase upon excitation^30^. Further, metabolites within the model were constrained based on our bat heart metabolomics analysis results. The uptake of glucose, ketone bodies, amino acids and lipids was not restricted. The GUROBI LP solver was used to find the solution to the FBA problems^31^.

### 2.5. Statistical Analysis

Statistical analysis was conducted using GraphPad Prism software (v.10.40.1) and R studio (v.2024.09.0). Differences between groups were considered significant at p<0.05. Sample sizes were not predetermined based on statistical power calculations. Formal randomizations of mouse or bat experiments were not used. Data were tested for normal distribution and similar variance among treatments using the Shapiro-Wilk tests. Statistical significance was calculated by multiple unpaired t-tests and comparisons analysis using a false discovery rate < 5% by the two-step method of Benjamini, Krieger, and Yekutieli.

## 3. RESULTS

The analysis of cardiac metabolomics data revealed that Egyptian fruit bats have a distinct cardiometabolic profile compared to C57/BL6J mice (PCA analysis plot, **Figure 1B**). Specifically, the abundance of 43 profiled metabolites significantly differed between bats and mice (**Figure 1C**). Notably, cardiac energy reserves were depleted in bat hearts, as evidenced by reduced creatine and undetectable glycogen (**Figure 1D**). Consistent with these findings, we found increased AMP levels in bat hearts (3.5-fold vs. mouse, **Figure 1E**), while NAD(P)^+^ redox equivalents were reduced. In the bat heart, glycolytic and Krebs cycle intermediates (**Figure 1F**) and amino acids (**Figure 1H**) were reduced compared to mice. Only constituents of lipid metabolism (**Figure 1G**) were universally increased (i.e., acetate, choline, phosphocholine, and carnitine), indicating an increased contribution of lipids to cardiac metabolism in bats. Correlation analysis of metabolic intermediates (**Supplementary Figure 1**) revealed co-regulation of ketogenic amino acids entering the Krebs cycle through acetyl-CoA. Increased ketone body metabolism spares protein degradation and further supports lipid oxidation. Together, our metabolomics findings suggest metabolic adaptation in the bats characterized by increased oxidation of lipids to meet cardiac energy demands.

Our systems biology approach revealed that bat hearts show a manifold increase in cardiac metabolic flux of oxidative phosphorylation, β-oxidation of long-chain fatty acids, glutamate, pyruvate, Krebs cycle (α-ketoglutarate), and total adenine nucleotide pool (AMP, ADP, ATP; **Figure 1A**). Furthermore, bat hearts were characterised by the markedly elevated flux of fructose, ketone body metabolism, and reduction in amino acid catabolic flux exchange (**Figure 1B** and **1C**).

Comparison of CardioNet-simulated predicted flux distributions from different activity states in bats (night, roost, and flight) show different cardiac metabolic profiles (**Supplementary Figure 2B**). Predicted myocardial oxygen consumption increased with activity status (**Figure 2C**), with flight showing the highest consumption rate, followed by night and roost activity. These differences were also reflected in the consumption of energy-providing nutrients. The glucose utilisation is predicted to be low in all three activity states (**Figure 2D**). The uptake and oxidation of linoleate (FA18:2) is highest during flight, while β-hydroxybutyrate contribution increases significantly during roost and night activity. This difference was also reflected at the level of the mitochondrial electron transport chain fluxes (**Figure 2E**), where complex I and V fluxes increase significantly during flight and are lowest during roost activity. Thus, fatty acids are major energy-providing substrates, and ketone bodies further complement ATP provision during roost and night activities.

**Figure 2.**
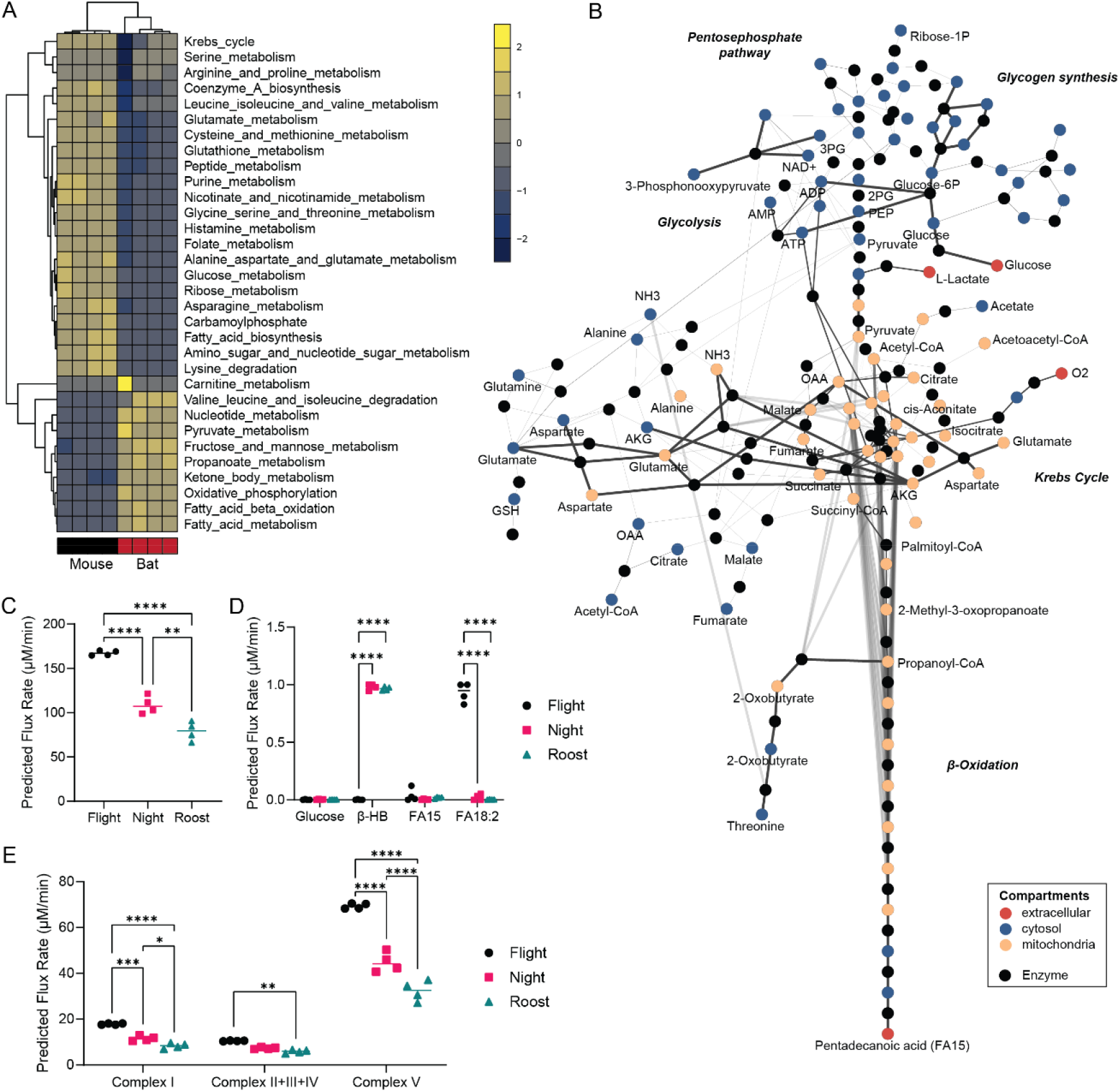
CardioNet analysis reveals increased lipid oxidation flux in bats. (**A**) Unsupervised hierarchical clustering of estimated z-scored flux rate changes reveals metabolic adaptation in bats vs mice. Z-scores were calculated to visualise how many standard deviations an estimated flux rate is away from the mean across all experimental groups. The z-score describes the distance from the mean for a given flux rate as a function of the standard deviation. The colour scale indicates the degree to which estimated flux rate changes are predicted to be respectively lower or higher bat vs mouse. (**B**) CardioNet *in silico*metabolic >5-fold flux changes bat versus C57/BL6J. The graph denotes the estimated flux distribution. The coloured nodes represent metabolites assigned to five compartments: extracellular space, cytosol, mitochondria, microsome, and lysosome. The black nodes indicate reactions; two reactions are linked by a directed edge, indicating the reaction flux. The line thickness of each edge is proportional to the predicted flux rate change. (**C**) Predicted myocardial oxygen consumption in simulations of bat flight, night activity and roost activity (n=4/group). (**D**) Predicted substrate uptakes for glucose, β-hydroxybutrate (β-HB) and fatty acids (FA15, FA18:2) in simulations of bat flight, night activity and roost activity (n=4/group). (**E**) Predicted flux rates for oxidative phosphorylation and electron transport chain in simulations of bat flight, night activity and roost activity (n=4/group).

## 4. DISCUSSION

Our study provides the first integrative metabolomic and flux balance analysis of Egyptian fruit bat (*Rousettus aegyptiacus*) myocardium, revealing distinct metabolic signatures compared to the widely used murine model (C57/BL6). By combining untargeted and targeted metabolomics with *in silico*modeling constrained by empirical data, we identify key pathways, particularly long-chain fatty acid and ketone body oxidation, that support the extreme energetic demands of the bat heart while minimizing reliance on carbohydrate reserves.

Classic physiological studies established that bats maintain exceptional cardiac performance during energetically costly flight^14,32^. Our findings align with this literature while extending it by identifying metabolic pathways that may underlie this performance. Our data suggest that, in bat hearts, glycogen stores are depleted and that lipid oxidation is the dominant energy-providing substrate. In the heart, glycogen reserves are normally utilised during periods of high physical activity^33^ or a means of adaptation to extreme physiological environments, such as was recently observed in naked mole rats^23^. High AMP concentration signals a highly catabolic cellular state and allosterically inhibits phosphofructokinase in glycolysis, which supports replenishing glycogen pools^29^. Our cardiac metabolic profile results align with isotopic tracer studies of systemic metabolism demonstrating that *R. aegyptiacus*rapidly and directly oxidizes exogenously supplied substrates to fuel both rest and flight. ^34^ Simulation of activity states revealed that the bat heart shifts from predominately oxidizing fatty acids (FA18:2) to ketone bodies (β-hydroxybutyrate) during lower activity states at night and during roosting. Interestingly, myocardial ketone body and branched-chain amino acid oxidation exceed glucose oxidation *in silico*, which requires further experimental validation. While the frugivorous diet of *R. aegyptiacus*is relatively protein-poor, our model predictions of greater amino acid oxidation in cardiac tissue likely reflect the endogenous catabolism of intracellular amino acid pools, rather than a reliance on dietary amino acids *per se*.

Our previous work^21^ in multiple non-cardiac tissues demonstrated positive selection in carbohydrate metabolism genes across nectar- and fruit-feeding bats, consistent with a high capacity for carbohydrate flux. In our study, the *R. aegyptiacus*heart showed high predicted flux through oxidative pathways, including fatty acid oxidation, at the time of sampling. These findings are not mutually exclusive: while the bat heart may retain the capacity for rapid carbohydrate oxidation under fed or exercise conditions, cardiac energy supply during rest or fasting may rely predominantly on fatty acid oxidation. *R. aegyptiacus*has been shown to exhibit dramatic postprandial hyperglycemia, which would affect myocardial exogenous substrate supply^18,35,36^. Our tissue samples were obtained opportunistically, and thus could not control for feeding status or recent activity. The absence of detectable glycogen and elevated fatty acid oxidation fluxes in our data likely reflect the metabolic state at the time of sampling. Future studies combining plasma and tissue metabolomics under controlled fasted, fed, and exercise conditions will be crucial to capture the full dynamic range of cardiac fuel utilization.

A key advance of this study is integrating metabolite concentrations with flux balance analysis (CardioNet) to estimate pathway-level metabolic activity rather than relying solely on static metabolite abundances. This approach enables generation of mechanistic hypotheses; for example, that fatty acid and ketone oxidation dominate basal myocardial energy supply in *R. aegyptiacus*. Nevertheless, we acknowledge that flux predictions require experimental validation, ideally via isotope tracing or perfused heart studies under controlled workloads and nutritional states.

We explicitly acknowledge that our study has several limitations. We also recognize the discussions surrounding limitations of two-species comparisons^37^ and therefore interpret our results as exploratory. The C57BL/6J mouse serves here as a physiologically tractable reference rather than a phylogenetic control, allowing us to highlight pathways potentially linked to the bat heart’s exceptional energetic resilience. Broader comparative studies across multiple bat and non-bat species will be essential to distinguish lineage-specific adaptations from general mammalian variation. We note that comparative metabolic research in non-model mammals, particularly bats, is inherently constrained by ethical and conservation considerations. Indeed all bats are slowly reproducing, and many species are protected by law thus opportunities to access cardiac tissue are exceedingly rare. While our sample size is modest, the data provide a unique and otherwise unattainable insight into the metabolic physiology of this species.The metabolite profiles were consistent across individuals, supporting the robustness of main findings. Two-species design precludes broad evolutionary generalization and results should be viewed as exploratory, motivating larger comparative studies. Our study captures a single metabolic state, as feeding status and activity levels were uncontrolled, limiting interpretation of dynamic metabolic flexibility. While CardioNet was originally parameterized in model species, our analysis was constrained by *R. aegyptiacus*-specific metabolomic data, ensuring that flux predictions reflect measured myocardial states. We present these results as hypothesis-generating rather than definitive, and we provide all simulation code to facilitate future refinement using species-specific telemetry and metabolic data.

Despite limitations, our study offers rare and valuable insights into bat cardiac metabolism, highlighting a shift toward oxidative substrates and minimal cytosolic energy reserves. We propose that metabolic flexibility switching between carbohydrate, lipid, and ketone oxidation may underpin the bat heart’s ability to sustain extreme energetic demands without pathological remodeling. Future work should prioritize controlled feeding and exercise experiments with paired plasma-tissue metabolomics, isotope tracing to validate flux predictions and multi-species comparisons to dissect ecological vs. phylogenetic influences on cardiac metabolism. Such studies will clarify whether the pathways highlighted here represent general principles of chiropteran cardiac physiology or unique adaptations of *R. aegyptiacus*.

## CRediT authorship contribution statement

**A.K**.: Writing – review & editing, Writing – original draft, Validation, Methodology, Investigation, Formal analysis. **F.C**.: Writing – review & editing, Investigation, Formal analysis. **R.D**.: Writing – review & editing, Investigation, Formal analysis. **K.K**.: Writing – review & editing, Investigation, Formal analysis. **M.Y**.: Writing – review & editing, Investigation, Formal analysis. **S.J.R**.: Writing – review & editing, Investigation, Formal analysis. **D.A**.: Writing – review & editing, Investigation, Writing – original draft, Supervision, Funding acquisition, Formal analysis, Conceptualisation.

## Data statement

All the data acquired are presented in the figures.

## Acknowledgements

We are grateful to Dr Mads Bertelsen (Copenhagen Zoo, Denmark) for facilitating access and helping with bat tissue collection. We acknowledge Barts Cancer Institute metabolic flux analysis facility (Dr Katiuscia Bianchi and Valle Moralles) for LC-MS/MS sample analysis and Dr Nasima Kanwal (School of Physical and Chemical Sciences Queen Mary University of London) for technical assistance with ^1^H NMR spectroscopy. Funding: This work was supported by the National Institutes of Health (NIH) (R00-HL-141702 and R01-HL177461 to A.K.), Barts Charity (G-002145DA) and Wellcome Trust (221604/Z/20/Z DA). The metabolic flux analysis facility of the Barts Faculty of Medicine and Dentistry was created with the support of the Barts and the London Charity (grant number MGU0401).

## Supplementary Figures

**Supplementary Figure 1.**
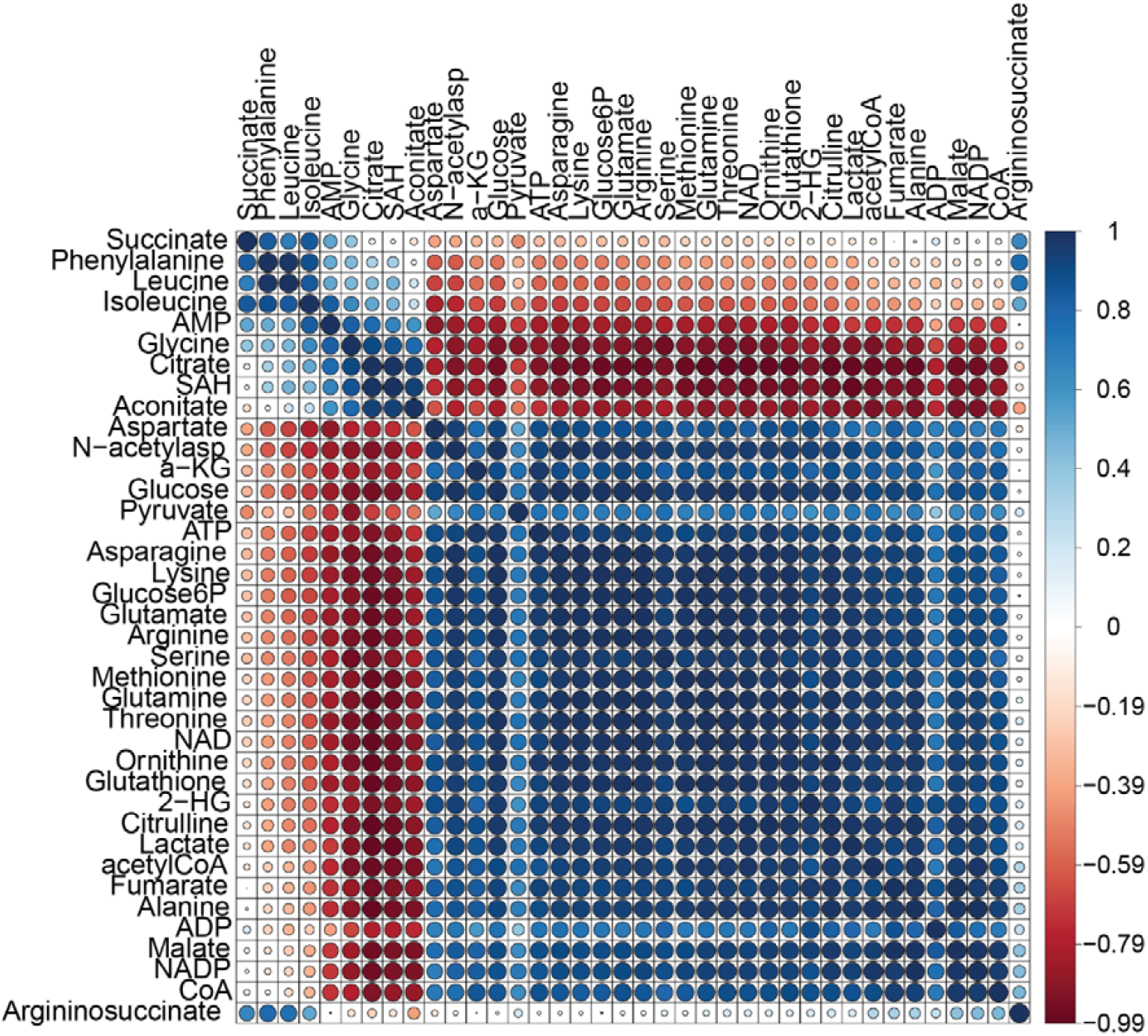
Correlation analysis of bat metabolome. Metabolite abundances from NMR- and LC-MS/MS-based metabolomics were analysed using Spearman correlation to reveal interactions between intermediates. Positively and negatively correlating intermediates are indicated by blue and red colour coding, respectively. Circle sizes indicate calculated absolute spearman coefficient ranging from 0 to 1.

**Supplementary Figure 2.**
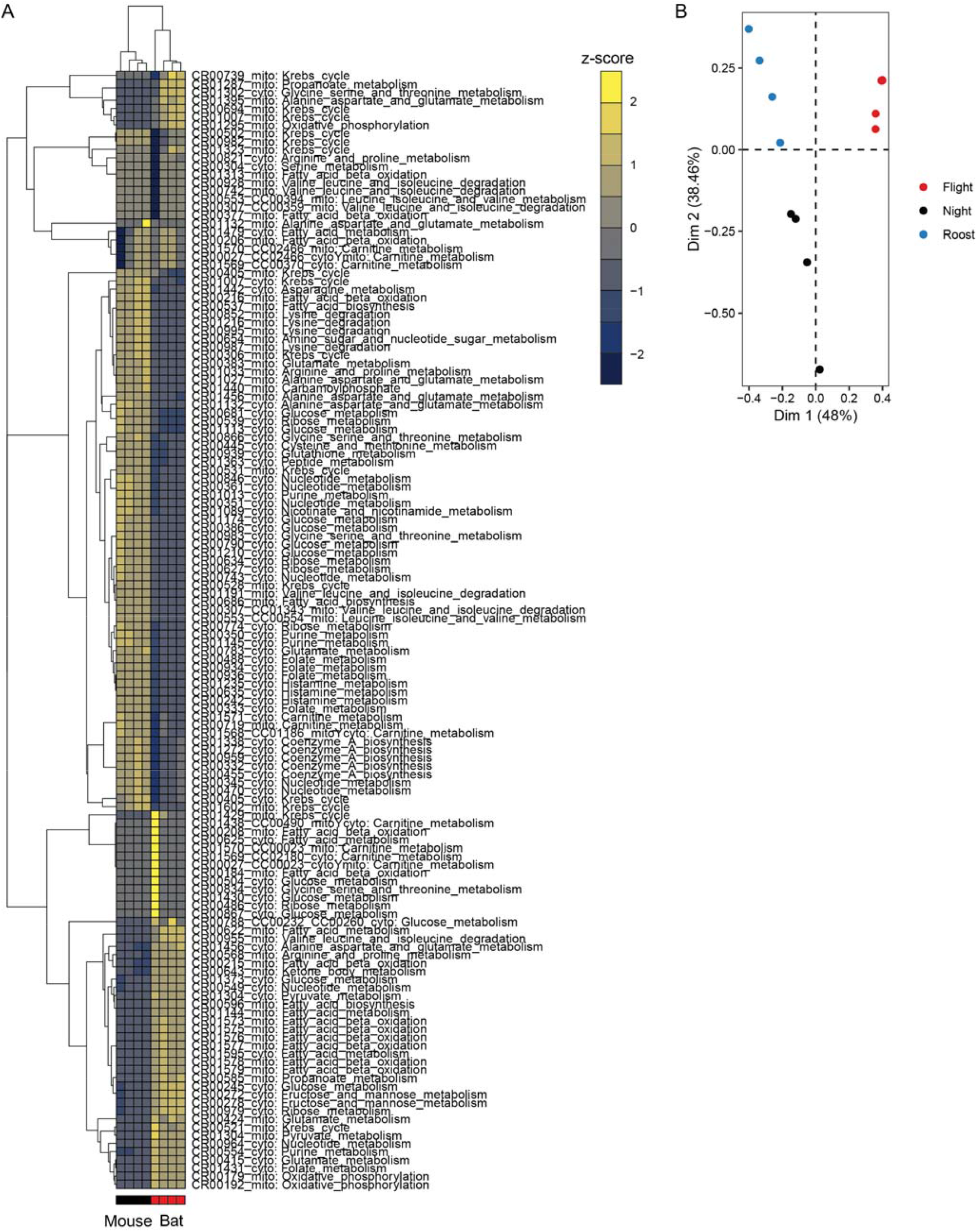
*In silico*modelling of bat metabolism. (**A**) Unsupervised hierarchical cluster analysis and heatmap of significantly altered metabolic fluxes in bat and mouse simulations using CardioNet. Flux rates were z-score normalised. n=4/species. (**B**) Principal component analysis of simulated activity states: flight, night and roost.

